# Metagenomic Discovery of CRISPR-Associated Transposons

**DOI:** 10.1101/2021.08.16.456562

**Authors:** James R. Rybarski, Kuang Hu, Alexis M. Hill, Claus O. Wilke, Ilya J. Finkelstein

## Abstract

CRISPR-associated transposons (CASTs) co-opt Cas genes for RNA-guided transposition. CASTs are exceedingly rare in genomic databases; recent surveys have reported Tn7-like transposons that co-opt Type I-F, I-B, and V-K CRISPR effectors. Here, we expand the diversity of reported CAST systems via a bioinformatic search of metagenomic databases. We discover new architectures for all known CASTs, including novel arrangements of the Cascade effectors, new self-targeting modalities, and minimal V-K systems. We also describe new families of CASTs that have co-opted the Type I-C and Type IV CRISPR-Cas systems. Our search for non-Tn7 CASTs identifies putative candidates that co-opt Cas12a for horizontal gene transfer. These new systems shed light on how CRISPR systems have co-evolved with transposases and expand the programmable gene editing toolkit.

## Introduction

CRISPR-associated transposons (CASTs) are transposons that have delegated their insertion site selection to a nuclease-deficient CRISPR-Cas system. All currently-known CASTs derive from Tn7-like transposons and retain the core transposition genes TnsB and TnsC but dispense with TnsD and TnsE, which mediate target selection (1, 2). Tn7 transposons site-specifically insert themselves at a single chromosomal locus (the attachment or *att* site) via the TnsD/TniQ family of DNA-binding proteins, while TnsE promotes horizontal gene transfer onto mobile genetic elements. In contrast, Class 1 CASTs replace TnsD and TnsE with a crRNA-guided TniQ-Cascade effector complex (3–6). These CASTs can use the TniQ-Cascade complexes for both vertical and horizontal gene transfer (5). One notable exception is a family of Type I-B CASTs that retains TnsD for vertical transmission but co-opts TniQ-Cascade for horizontal transmission (7). Similarly, Class 2 CASTs use the Cas12k effector to transpose to the attachment (*att*) sites or to mobile genetic elements (8, 9). CASTs also dispense with the spacer acquisition and DNA interference genes found in traditional CRISPR-Cas operons (2). In short, these systems have merged the core transposition activities with crRNA-guided DNA targeting.

CASTs are exceedingly rare; only three sub-families of Tn7-associated CASTs have been reported bioinformatically and experimentally (2, 5, 7, 9, 10). These studies have identified that many, but not all, CASTs encode a self-targeting spacer flanked by atypical (privileged) direct repeats. However, the prevalence of such atypical repeats, the diversity of self-targeting strategies, and the molecular mechanisms of why CASTs have evolved these repeats remain unresolved. Moreover, all CASTs that have been identified to-date have a minimal CRISPR array with as few as two spacers. These systems are also missing the Cas1-Cas2 adaptation machinery, raising the question of how CASTs target other mobile genetic elements for horizontal gene transfer. Another open question is whether non-Tn7 transposons have adapted CRISPR-Cas systems to mobilize their genetic information. We reasoned that an expanded catalog of CASTs may shed light on many of these unresolved questions.

Here, we have systematically surveyed CASTs across metagenomic databases using a custom-built computational pipeline that identifies both Tn7- and non-Tn7 CASTs. Using this pipeline, we have identified unique architectures for Type I-B, I-F, and V CASTs. Type I-F CASTs show the greatest diversity in Cas genes, including TniQ-Cas8/5 fusions, split Cas7s, and even split Cas5 genes. Some I-F CASTs appear likely to assemble a Cascade around a short crRNA for self-targeting from a non-canonical spacer. Type I-B CASTs frequently encode two TniQ/TnsD homologs, one of which is used for self-targeting via a crRNA-independent mechanism. Remarkably, we have also found I-B systems that encode two TniQ homologs and a self-targeting crRNA, suggesting additional unexplored targeting mechanisms. In addition, we have observed new Type I-C and Type IV-family Tn7-like CASTs with unique gene architectures. Both of these sub-families lack canonical CRISPR arrays, suggesting that CASTs use distal CRISPR arrays, perhaps from active CRISPR-Cas systems, for horizontal gene transfer. We have identified multiple self-insertions and gene loss in Type V systems, indicating that target immunity—a mechanism that prevents transposons from multiple self-insertions at an attachment site—is frequently weakened. Finally, we have found a set of Cas12a-associated Rpn-family transposases that may participate in crRNA-guided horizontal gene transfer. We anticipate that these findings will shed additional light on how CASTs have co-opted CRISPR-Cas systems and further expand the precision gene editing toolbox.

## Results

### A bioinformatic CRISPR-associated transposon (CAST) discovery pipeline

We developed a bioinformatics pipeline that first searches metagenomic contigs for transposases using protein BLAST (BLASTP) and a curated transposase database with a permissive e-value threshold of 10^−3^ (11) (**Figure 1A**). All possible open reading frames (ORFs) in a 25 kilobasepair (kbp) neighborhood up- and downstream of each putative transposase are translated and searched with BLASTP using a second curated database of all Cas proteins. Contigs without Cas genes are not analyzed further. The remaining contigs contain both a transposase and at least one Cas gene. We identify CRISPR arrays in these contigs using a modified version of PILER-CR that can locate arrays with as few as two repeats (12). A final round of protein annotation searches for accessory transposase subunits (i.e., TnsC/D/E for the Tn7 family) and genetic elements that are near common attachment sites (13, 14). Finally, we filter the contigs by constraints (detailed in later sections) that are designed to isolate novel CASTs.

**Figure 1.**
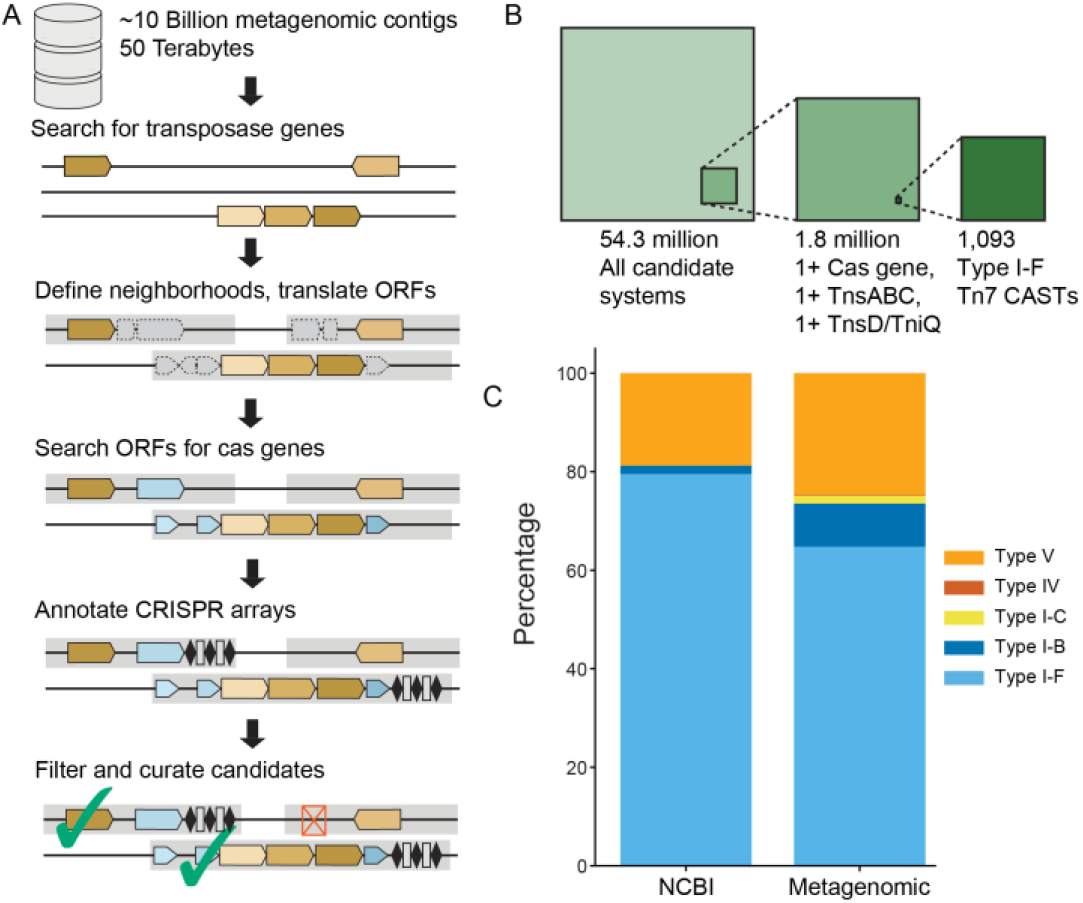
CRISPR-associated transposon detection and classification. **(A)** A bioinformatic pipeline for the discovery of CRISPR-associated transposons (CASTs). Brown: transposase genes; blue: cas genes; dotted: open reading frames; gray: gene neighborhoods. Neighborhoods satisfying initial search criteria are marked with a green check. Red “x” denotes a neighborhood that doesn’t match initial search criteria (e.g., no detected cas genes). **(B)** Summary of the stepwise filtering strategy to identify high-confidence Type I-F Tn7 CASTs. **(C)** Distribution of Tn7-associated CAST subtypes in the NCBI microbial genome and EMBL metagenomic databases.

To test this pipeline, we searched for all previously discovered CASTs in the National Center for Biotechnology Information (NCBI) repository of bacterial genomes (2, 15, 16). We downloaded 951,491 partial and complete bacterial genomes from the NCBI FTP server on May 5, 2021. Using these genomes as input, we identified regions with at least one Cas gene within a ∼25 kbp neighborhood of a transposase. We filtered for contigs that include either a Type I or Type V effector and at least one member of the TniQ/TnsD family of Tn7-associated proteins. As expected, we re-identified previously published systems, along with previously unannotated Type V systems. These results confirm that the bioinformatics pipeline is sufficiently sensitive to discover these rare CRISPR-transposon systems in large genomic databases.

Next, we searched the EMBL-EBI repository of metagenomic sequencing reads for novel CASTs (17). This repository is the largest collection of sequenced DNA from diverse microbiomes, aquatic, man-made, and soil environments. We downloaded >1 petabyte of non-16S rRNA reads (**Figure 1B**, see **Methods**). These reads were assembled into ∼10 billion high-quality contigs (∼30 terabytes). Contigs were annotated for transposases and cas genes as described above. Approximately 54 million contigs (∼150 gigabytes) met the twin criteria of having at least one transposase and one cas gene. After de-duplication of nearly identical contigs, we searched for putative CAST systems (Figure 1B). Using Class 1 Tn7-associated CASTs as an example, we filtered for contigs that included at least one cas gene, tnsD/tniQ, and had at least one of the core Tn7 genes: tnsA, tnsB, or tnsC (∼1.8M contigs). These systems were additionally filtered by selecting only systems where two of the Tn7 genes were less than 1500 bp apart, and where three of the core Class 1 cas genes (cas5, cas6, cas7, and cas8) were also less than 1500 bp apart (1167 systems). Of these, 1093 systems were identified as being Type I-F CASTs. The remaining high-confidence systems included Type I-B, I-C, IV, and V systems (Figure 1C). To increase the confidence of our annotations, and to annotate unknown ORFs, we re-BLASTed all possible reading frames against the UniProtKB/TrEMBL database of high-quality computationally annotated protein variants (18). We assigned high confidence systems to specific sub-types by re-BLASTing the cas genes against a database of subtype-specific effector proteins and by manually reviewing the operon architecture. After removing nearly redundant systems, we found 1476 high-confidence CRISPR-Tn7 CASTs. Notably, we detected founding members of the Type I-C and IV CASTs in the metagenomic but not NCBI database (see below). All of these systems were missing the interference (Cas3) and adaptation (Cas1/2) genes, in agreement with the ‘guns for hire’ hypothesis for how CRISPR-Cas systems have been co-opted for diverse cellular functions (19).

### Diversity of Type I-F CASTs

We identified 1093 non-redundant I-F sub-systems with the prototypical gene arrangement of TnsA-TnsB-TnsC separated from TniQ-Cas8/Cas5-Cas7-Cas6 by a large cargo region (**Figure 2A**). Tn7 cargo genes are unrelated to the transposition mechanism, and often include antibiotic resistance genes (20). Type I-F3a systems, defined as using the conserved genes guaC or yciA as their attachment site, comprise ∼61% of all I-F CASTs (**Figure 2B**). I-F3b systems, which use the rsmJ or ffs attachment site, comprise ∼34% of I-F CASTs (21). The remaining 5% of I-F systems form a distinct group, termed I-F3c, with a unique attachment site and self-targeting mechanism (see below).

**Figure 2.**
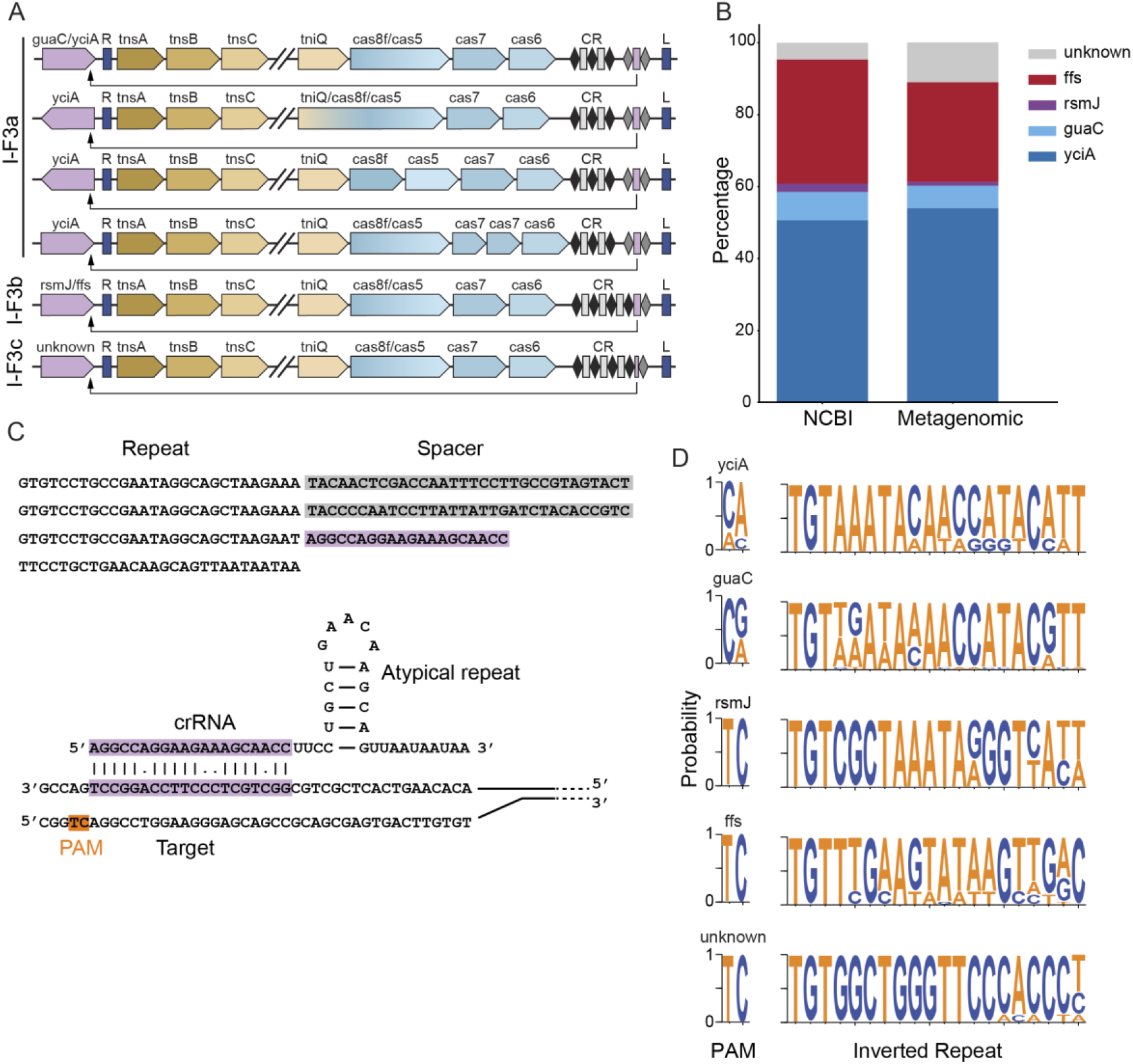
Summary of Type I-F CASTs. **(A)** Gene architectures of Type I-F3a, I-F3b, and I-F3c systems. Unique gene architectures include TniQ-Cas8 fusions, split Cas8 and Cas5, and dual Cas7 systems. Purple: attachment site; blue: left (L) and right (R) transposon ends. Black diamonds: canonical direct repeats; gray diamonds: atypical direct repeats. Rectangles: protospacers; purple rectangle: self-targeting protospacer. Arrow indicates the target site. Slanted gapped lines indicate elided cargo regions. **(B)** The distribution of attachment site genes in the NCBI and the metagenomic databases. **(C)** (top) Sequence of a CRISPR array with a short, atypical spacer (purple) that may assemble a mini-Cascade. (bottom) Schematic of an atypical crRNA and its target DNA sequence. **(D)** Weblogos of the PAM and right inverted repeat adjacent to each attachment site. The TnsB binding site and the self-targeting PAMs are conserved within sub-systems

The most common gene arrangement in our dataset for all three subtypes encodes the TnsA-C proteins in one operon, and the TniQ and Cas proteins in a second operon that is adjacent to the CRISPR repeats. A large cargo spanning ∼10-20 kbp either separates these operons or is present downstream of the cas genes. The stoichiometry of the Cascade effector has been previously reported to be (Cas6)_1_ : (Cas7)_6_: (Cas8/Cas5)_1_ : crRNA_1_ : TniQ_2_, based on cryo-electron microscopy of Type I-F CAST complexes (3, 4, 6, 22). In these studies, TniQ interacts with Cas7 and is structurally distant from Cas8/Cas5. However, in two of our new I-F3a systems, TniQ is expressed as an N-terminal fusion with Cas8/Cas5. Four distinct I-F3a systems also have a split Cas7, and in one system, both the Cas5 and Cas7 proteins are split into two distinct polypeptides.

All I-F3 systems that we identified appear to use a crRNA-guided self-targeting mechanism that directs Cascade near the attachment site (21). The self-targeting crRNA is either in the leader-distal position of the CRISPR array or 80-85 nt away from the CRISPR array, as reported previously (21). These self-targeting crRNAs are flanked by an atypical direct repeat that has several substitutions relative to the direct repeats within the CRISPR array. I-F3c CASTs attach upstream of a protein of unknown function that encodes seven putative transmembrane regions (see **Methods**). This attachment site has not been previously reported for any Tn7-family transposon. To determine how Type I-F3c systems use crRNA-guided transposition, we aligned the region around the CRISPR array with the sequence 500 bp upstream of tnsA. This identified a short 20 bp sequence immediately after the final canonical CRISPR repeat that matched the region approximately 64 bp upstream of the transposon end. This short, atypical spacer is followed by an atypical repeat (**Figure 2C**), akin to the I-F3a and I-F3b systems.

Type I-F systems recognize a dinucleotide protospacer adjacent motif (PAM) (23). Our analysis of the self-targeting PAMs highlighted that they vary with the attachment site and CAST sub-family (**Figure 2D**). Next, we analyzed the sequence composition of the inverted repeats that span Tn7. The right inverted repeat starts with a universally conserved 5’-TGT that is recognized by the essential TnsB recombinase (24). The rest of this repeat varies but is most similar between CASTs that have the same attachment site (**Figure 2D**). These results further confirm that I-F3c systems cluster into a distinct CAST sub-type.

### Type I-B CASTs encode multiple integration mechanisms

We found four families of Type I-B CASTs that lack interference and adaptation genes. These systems either encode a single TniQ or dual TniQs of unequal length (**Figure 3A**). Systems with dual TniQs comprise 79% of all identified systems. The most common Type I-B system (I-B1) encoded TniQ_1_ between TnsC and a Cas gene and TniQ_2_ is on the distal end of the CRISPR array (**Figure 3A, top and Figure 3B**). Systems with a single TniQ (I-B2 and I-B3) had two distinct gene architectures and self-targeting modalities. In I-B2 systems, where TniQ was sandwiched between TnsC and Cas6, we identified a self-targeting spacer that was complementary to a region downstream of glmS (**Figure 3A, middle**). However, we did not find any self-targeting spacers in I-B3 systems, where TniQ was between the CRISPR array and cargo genes. In addition, this TniQ was nearly double the length of the shorter TniQs found between TnsC and Cas6 and bore a strong resemblance to the TnsD encoded in canonical Tn7 systems. Notably, we also found single-TniQ systems that encode a TnsD-like TniQ but also lack self-targeting spacers.

**Figure 3.**
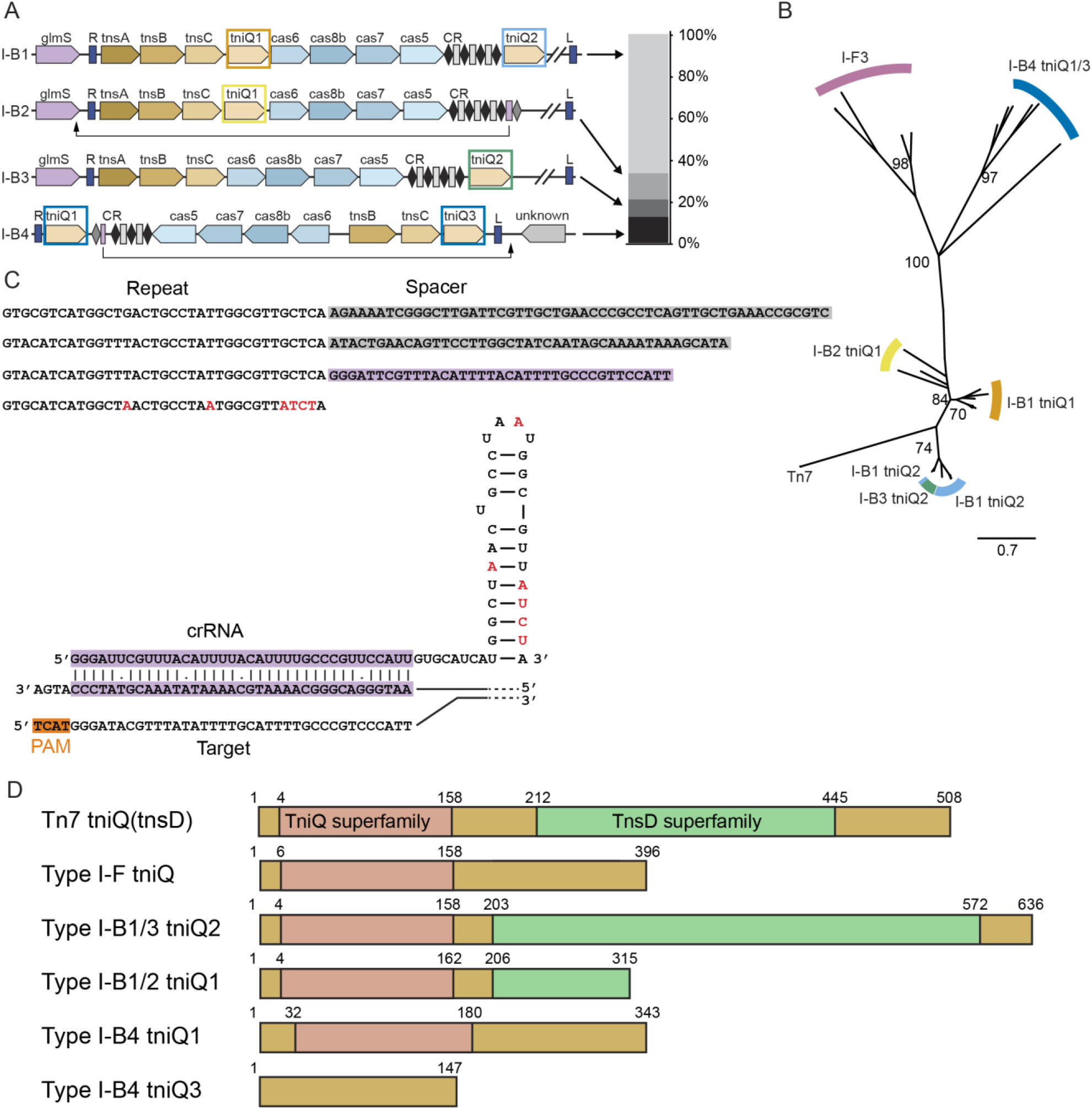
Analysis of Type I-B CASTs. **(A)** (left) Gene architectures of Type I-B systems. Systems can dispense with either the first or the second tniQ, suggesting alternative targeting lifestyles. Type I-B4 systems have a unique architecture that most resembles Type V CASTs. Colored rectangles correspond to phylogenetic groups in panel B. (right) The distribution of Type I-B sub-systems in the metagenomic database. **(B)** Phylogenetic tree with tniQ variants from Type I-B and I-F CASTs, as well as from the canonical Tn7 transposon. Type I-B tniQ1 is most similar to tniQ from Type I-F CASTs, whereas tniQ2 is closely related to canonical Tn7 tnsD. Values at branch points are bootstrap support percentages. **(C)** (top) Sequence of a Type I-B4 CRISPR array with a short, atypical spacer. (bottom) Schematic of an atypical crRNA basepaired with a target DNA sequence. Red bases are those that differ from the canonical repeat sequence. **(D)** Domain maps of TniQ proteins. Regions homologous to the TniQ superfamily and the TnsD superfamily are indicated in pink and light green, respectively. The Type I-B4 system encodes the shortest TniQ variant.

We identified an atypical Type I-B CAST (I-B4) that had a unique gene architecture and self-targeting mechanism (bottom row, **Figure 3A**). This system encodes TnsB and TnsC but lacks the TnsA gene, akin to Type V systems (8, 9). Two TniQ homologs of unequal length are immediately adjacent to the inverted repeats but distal from the Cas operon. TniQ_1_ is sandwiched between the right transposon end and a short CRISPR array; TniQ_3_ is only ∼450 bp long and is located between TnsC and the left transposon end. This short TniQ_3_ can be aligned against the N-terminus of traditional CAST-associated TniQs (**Figure 3D**). Notably, this is the only dual-TniQ CAST that encodes a self-targeting spacer with near-perfect complementarity to a region of DNA just outside the left transposon end. The attachment site is adjacent to a gene of unknown function near the left transposon end (**Figure 3A**), akin to the attachment sites in Type V CASTs. The self-targeting spacer is flanked by an atypical direct repeat and is also 6 to 23 bp shorter than the other spacers in the CRISPR array (**Figure 3C**). Based on these findings, and our observation of Type I-F CASTs with short self-targeting spacers, we propose that Type I-B systems can also assemble self-targeting mini-Cascades.

To better understand the roles of multiple TniQs in Type I-B CASTs, we constructed a phylogenetic tree of Type I-B, I-F3a, and I-F3b TniQs, along with TnsD from the canonical Tn7 transposon (**Figure 3B**). Compared with the cluster of TniQ_2_, the short TniQ_1_ from Type I-B1 and B2 systems are closer to TniQ from other CASTs, while the Type I-B1 TniQ_2_ clusters with canonical Tn7 TnsD. The close similarity between TniQ_2_ and Tn7 tnsD, along with the apparent lack of self-targeting in most dual TniQ systems, suggests that TniQ_2_ serves the same role as TnsD, namely that it is a sequence-specific DNA-binding protein that directs transposition downstream of glmS. TniQ_1_, in turn, forms a complex with Cascade to guide TnsABC to a crRNA-directed target. While this work was in preparation, an experimental study demonstrated that I-B CASTs with two tniQ genes use two separate pathways for target selection (7).

### Novel Type I-C and IV CASTs from metagenomic sources

Our analysis of the metagenomic contigs revealed a Type IV CAST (**Figure 4A**). Type IV systems are primarily encoded by plasmid-like elements to mediate inter-plasmid conflicts (25, 26). Phylogenetic trees of Cas6 and Cas7 independently placed this CAST within the Type IV-A3 sub-family (**Figure 4B**). These systems frequently shed their CRISPR repeats, instead of using distal CRISPR arrays (25). Although we did not find any CRISPR repeats in this system using CRASS or PILER-CR, we detected a spacer-like DNA segment with strong complementarity to the C-terminus of glmS, the likely attachment site. This putative self-targeting spacer is adjacent to a hairpin that resembles the direct repeats in other Type IV CRISPR-Cas systems (**Figure 4A, bottom**). We conclude that this minimal spacer-repeat motif directs self-targeting by the Type IV system. Horizontal transfer may still occur via a distal CRISPR array, akin to the interference mechanism in other Type IV CRISPR-Cas systems (25).

**Figure 4.**
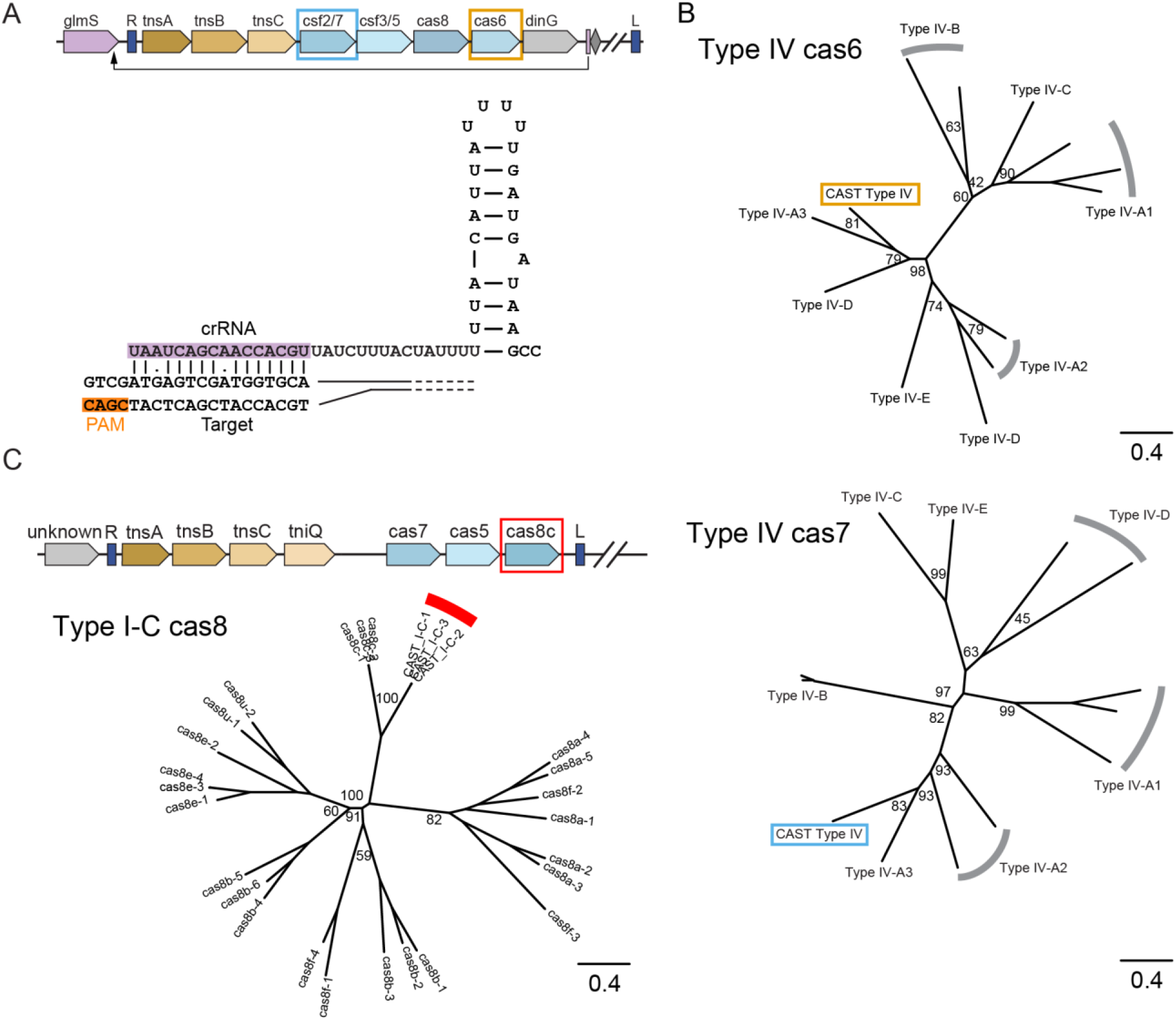
New Tn7 CASTs from metagenomic databases. (**A**) (top) Gene architecture of a Type IV CAST. This system lacks a CRISPR array but encodes a self-targeting spacer. Genes highlighted by colored rectangles correspond to genes in panel B. (bottom) Schematic of a short, self-targeting spacer basepaired with its target DNA sequence. **(B)** Phylogenetic trees of cas6 and cas7 indicate that the Type IV CAST most closely resembles Type IV-A3 CRISPR-Cas systems. Values at branch points are bootstrap support percentages. **(C)** (top) Gene architecture of Type I-C systems. We did not detect any CRISPR arrays or atypical self-targeting spacers. (bottom) A phylogenetic tree of cas8 confirms that this system is closely related to Type I-C Cascades. Values at branch points are bootstrap support percentages.

We found nine non-redundant Type I-C CASTs with Cas5, Cas7, and Cas8 downstream of TnsABC and TniQ (**Figure 4C**). TniQ is immediately adjacent to TnsABC rather than the Cas proteins. A phylogenetic analysis of Cas8 showed close similarity to Cas8c (**Figure 4C, right**). We did not detect a CRISPR array via CRASS or PILER-CR. We also did not detect any tRNA or common Tn7-associated attachment sites near either the left or right transposon ends, precluding a detailed analysis of the self-targeting mechanism. We cannot rule out that these systems use a minimal leader-spacer array that was below the threshold of our detection software. Alternatively, these systems may use a distal CRISPR array *in trans*. Further bioinformatic and experimental analyses will be required to delineate the mechanisms of self-targeting and horizontal transfer in these systems.

### Architectural diversity and self-targeting in Type V CASTs

Type V CASTs were likely formed when a Tn7-like transposon co-opted a Cas12 gene for RNA-guided DNA targeting (10, 15). Most Type V CASTs contain TnsB, TnsC, and TniQ at one end of the transposon with Cas12k, a small CRISPR array, and an atypical repeat-spacer on the other end. Cargo genes spanning 2 to 23 kb of additional DNA sequences are sandwiched between TniQ and Cas12k (**Figure 5A**). In contrast to the Class 1 systems described above, all metagenomic Type V systems lacked TnsA, consistent with the proposal that Cas12k was captured by a Tn5053-family transposon, which contains TnsB, TnsC, and TniQ homologs but also lacks TnsA (27). The tracrRNA is upstream of the canonical CRISPR array with homology to the atypical repeat. The crRNA usually has good homology to the target DNA, with 98% of systems containing one or zero mismatches in the first ten basepairs (**Figure 5B**). As previously observed (8), atypical spacers generally targeted tRNA genes immediately adjacent to the transposon. We also found one system that attaches 104 bp downstream of ArsS, an arsenosugar biosynthesis radical SAM protein. Analysis of the DNA upstream of the self-targeting spacer revealed 5’-TGGTA as the most common PAM, with some variability in the -5, -4, and -1 positions. (**Figure 5C)**. Experimental evidence for two CASTs showed a preference for a smaller 5’-GTN PAM (9). Whether the broader set of Cas12k proteins have more stringent PAM requirements will require experimental validation. Overall, this architecture and preference for tRNA attachment sites corroborate previous bioinformatic and experimental observations (7–9, 15).

**Figure 5.**
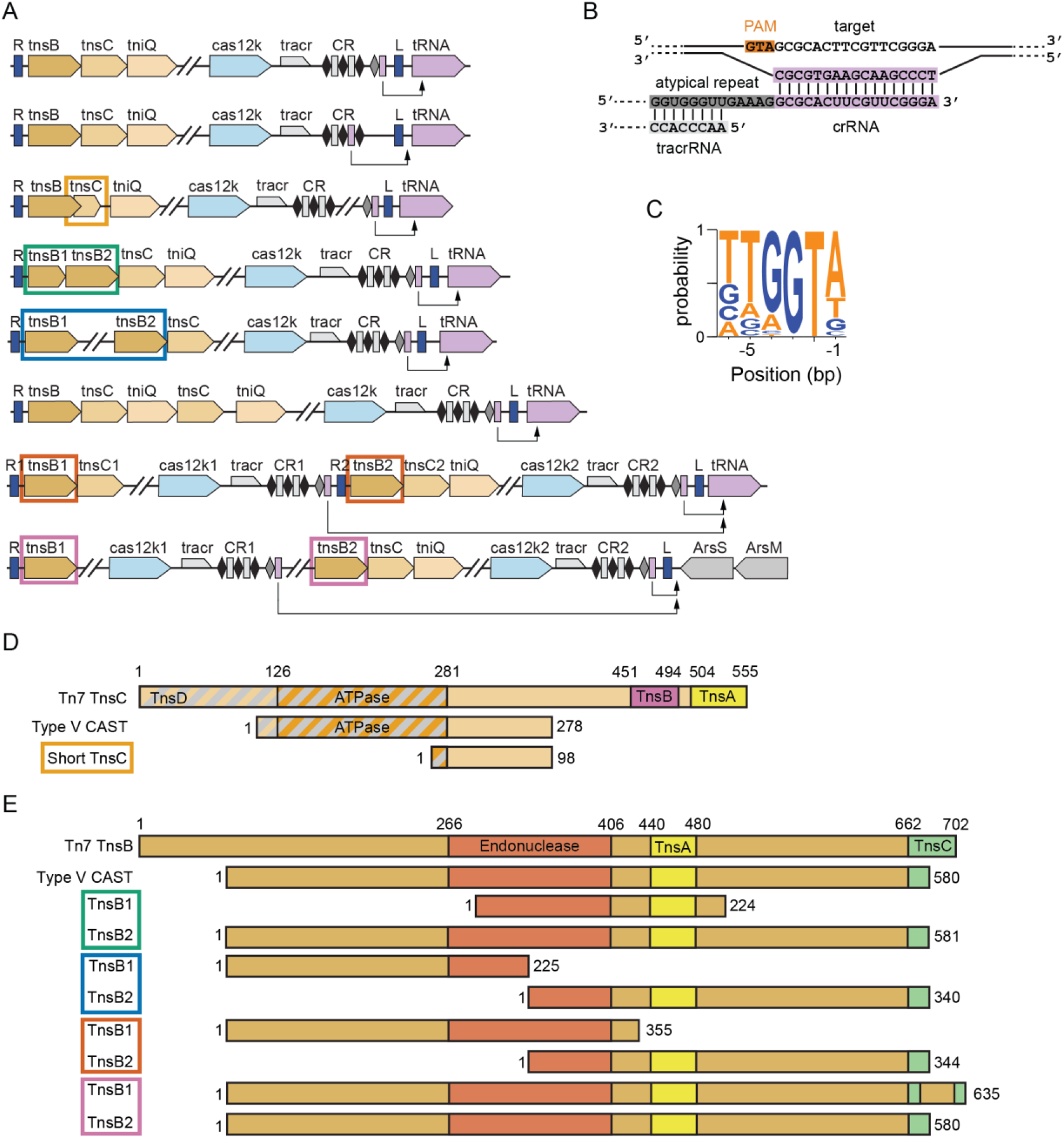
Analysis of Type V CASTs. (**A**) Gene architectures of Type V CASTs, including dual-insertion systems (bottom two rows). Colored rectangles around genes correspond to alignments in panels D and E. (**B**) Schematic of interactions between the target site DNA, a self-targeting crRNA, and a tracrRNA. (**C**) Weblogo of PAM sequences found adjacent to spacer targets. (**D**) Aligned domain maps of truncated TnsC variants. Gray diagonal stripes indicate TnsD-interacting region. Truncated TnsCs lack the TnsA- and TnsB-interacting domains but generally retain the ATPase domain and most of the TnsD-interacting domain. The shortest TnsC has also lost its ATPase domain. (**E**) Aligned domain maps of truncated TnsB variants. Type V CAST TnsB is shorter than Tn7 TnsB but contains the functionally annotated domains. In some dual TnsB systems, the first TnsB encodes the N-terminal region and the second encodes the C-terminal portion.

We also found Type V CASTs with unusual TnsC and TnsB arrangements. Notably, all Type V TnsC proteins lack the canonical TnsA- and TnsB-interacting domains and have partial truncations of the TniQ-interacting domain (28–30). The shortest CAST— only 6.6 kbp in total, including the cargo—encodes a 98 amino acid TnsC fragment whose sequence overlaps with TnsB by 115 bp (**Figure 5A, 5D**). In addition to losing the TniQ, TnsB, and TnsA interacting domains, this TnsC has also lost its ATPase domain (28, 30). We hypothesize that the minimal TnsC encodes uncharacterized TnsB- and TniQ-interaction motifs. Because of its compact organization, this CAST is also a prime candidate for gene editing applications.

Multiple CASTs split TnsB into separate ORFs that encode just the N- or C-terminus. Alternatively, a full-length TnsB is encoded next to a TnsB fragment containing most of the catalytic domain (**Figure 5E**). Strikingly, two unrelated systems encode the same N-terminal region of TnsB. We speculate that these split TnsBs form a heterodimeric TnsB_1_:TnsB_2_ transposition complex. These heterodimeric complexes retain the catalytic core while also maintaining the requisite TnsC interaction motifs via the longer TnsB subunit.

We also observed multiple contigs with two CASTs inserted at the same attachment site (**bottom rows, Figure 5A**). Tn7 transposons can prevent re-insertion at the attachment site by TnsB-mediated dissociation of TnsC from target DNA (31, 32). However, more distant Tn7-family transposases may still insert at a single attachment site, resulting in several transposons that are situated adjacent to each other (30). Consistent with this idea, the dual-insertion Type V CASTs have distinct cargos, unique gene architectures, and divergent Cas12k sequences. In one scenario, the tRNA-distal CAST encoded an N-terminal TnsB truncation and lost TniQ (7th row, **Figure 5A**). The tRNA-proximal CAST from the same organism encoded the C-terminal TnsB fragment and had a complete Type V-family TniQ. A second dual CAST system had lost both TnsC and TniQ from the tRNA-distal transposon (last row, **Figure 5A**). Both systems encode a self-targeting spacer and a full Cas12k gene, suggesting that they are both still active.

### Phylogenetic analysis of Tn7-CASTs

To clarify the evolutionary relationship between Tn7-CASTs, we built phylogenetic trees of the TnsB (**Supplemental Figure 1A**) and TnsC (**Supplemental Figure 1B**) proteins from all known CAST subtypes as well as Tn7-family transposons (see **Methods**). The phylogenetic relationships between sub-systems were nearly identical for both TnsB and TnsC, suggesting that these proteins are co-evolving as a system. We omitted TnsA from this analysis because Type I-B4 and all Type V Tn7-CASTs lack this gene. We confirmed that all metagenomic Type V-CASTs are phylogenetically closer to the Tn5053 transposon than Tn7. In contrast, Type I-B1-3, I-C, and IV CASTs are phylogenetically close to Tn7. Such limited evolutionary drift may suggest a relatively recent co-opting of this CRISPR-Cas system. Type I-B4 CASTs are a notable exception because these systems also lack TnsA and cluster closer to Tn5053 than to the reference Tn7. Type I-F CASTs are highly divergent from both Tn7 and Tn5053, with a large phylogenetic separation between the I-F3a and I-F3b sub-types. The diversity of Tn7-family transposases and CRISPR sub-systems that have co-evolved suggests that we will continue to find new CAST families as metagenomic databases expand and detection sensitivity increases.

### Non-Tn7 CASTs that co-opt Cas12a and Type I -E Cascade

While transposons other than Tn7 may have co-opted CRISPR-Cas systems for attachment site recognition, this has never been reported. To explore this possibility, we identified contigs that encode: (1) at least one non-Tn7 transposase gene (see **Methods**), (2) a CRISPR array containing at least two spacers, and (3) Class 1 or Class 2 DNA-binding effectors (i.e., Cas9, Cas12, Cas13, or any three of Cas5/6/7/8). We excluded contigs that encode interference (i.e., Cas3 or Cas10) or acquisition machinery (i.e., Cas1 or Cas2). Class 2 nucleases were additionally filtered by size to exclude truncated genes (see **Methods**). We prioritized Type II, V, and VI systems where the catalytic nuclease domain residues are mutated or deleted, as these enzymes cannot participate in adaptive immunity. We detected nuclease-inactivating mutations or deletions in 25% of Cas9 genes (in one or both nuclease domains), 8% of Cas12 genes, and none of the Cas13 genes.

We found 40 non-redundant examples of a nuclease-inactive Cas12a or a Type I-E Cascade near a putative recombination-<u>p</u>romoting <u>n</u>uclease/transposase (Rpn)-like protein (**Figure 6A**). Rpn family proteins were originally investigated because of their close homology to the catalytic domain of transposase_31 (33). These proteins contain a PDDEXK nuclease domain, first discovered in restriction endonucleases, but also observed in T7 TnsA and other diverse DNA-processing enzymes (34–36) (**Figure 6B**). *E. coli* RpnA promotes RecA-independent gene transfer in cells and is a Ca^2+^-stimulated DNA nuclease *in vitro* (33). The mechanism of how RpnA promotes horizontal gene transfer is unknown.

**Figure 6.**
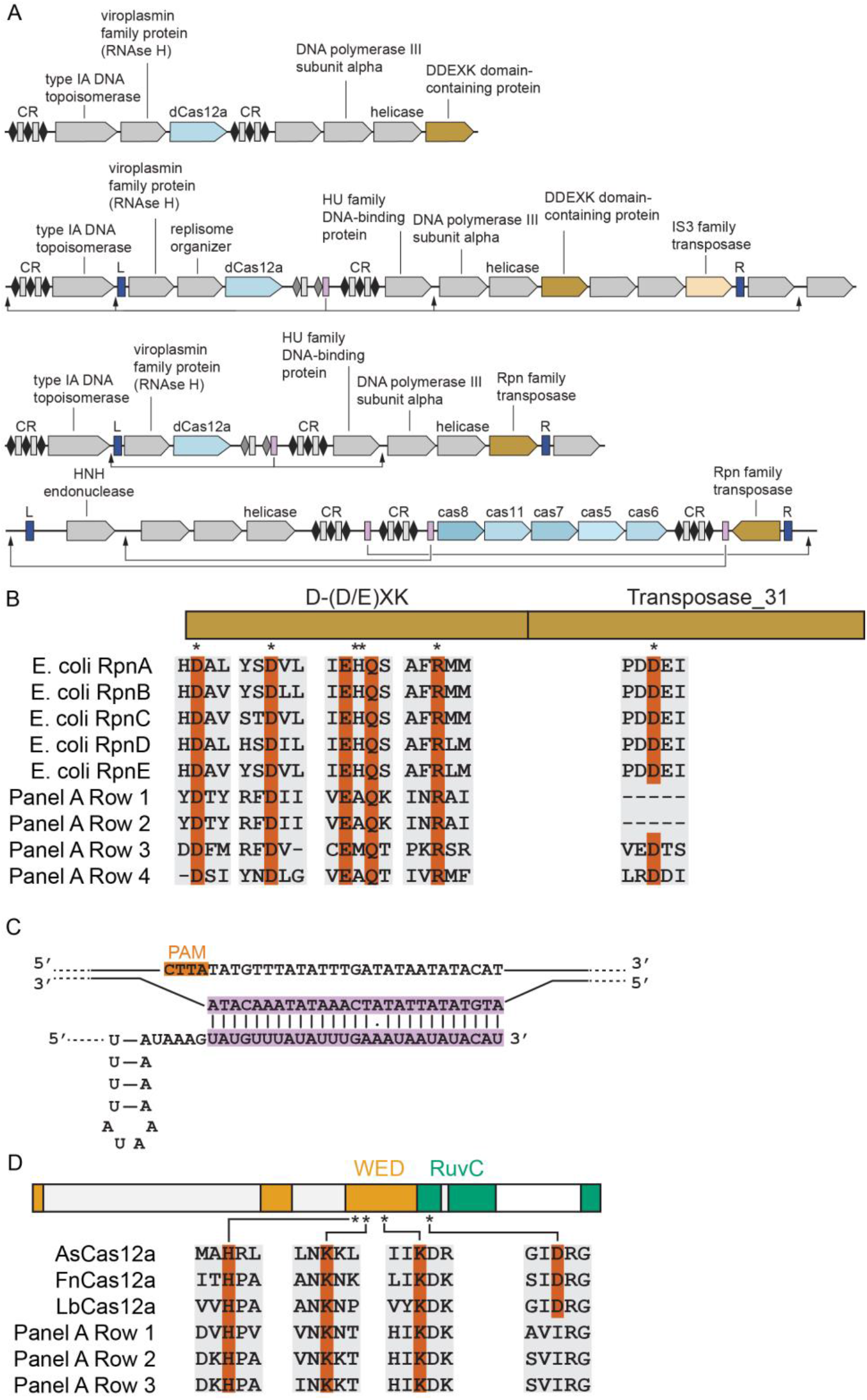
A family of putative non-Tn7 CASTs. **(A)** The defining features of this family of systems are an Rpn-family (PDDEXK domain-containing) nuclease/transposase near a nuclease-dead Cas12a or a Type I-E Cascade complex. The operon is enriched for nucleic-acid processing proteins. We also observed self-targeting spacers (magenta, black arrows) and short inverted repeats (blue) in some systems. **(B)** Multiple sequence alignment of Rpn proteins with the putative transposases from these systems. Residues critical for DNA cleavage in the PDDEXK domain are highlighted in red. The D165A mutant in RpnA more than doubles recombination *in vivo*; this aspartic acid is highlighted in red below the transposase_31 domain. **(C)** Schematic of an atypical self-targeting spacer and its DNA target. The PAM is highlighted. **(D)** Multiple sequence alignment of nuclease-active Cas12a and putative CAST Cas12as. Putative CAST Cas12as retain the conserved residues in the WED domain that are essential for crRNA processing, but lack an aspartic residue in the RuvC domain that is essential for DNA cleavage.

The genetic context around Cas12a in our systems is highly enriched with nucleic acid-interacting proteins, including a topoisomerase, an RNAse, a DNA polymerase subunit, and one or two helicases. Likewise, the Cascade system encodes a helicase and an HNH endonuclease. Three Cas12a systems encode an HU family DNA-binding protein, and one of those also contains a protein with homology to a phage replisome organizer (37). We detected three systems with putative atypical self-targeting spacers adjacent to a canonical CRISPR array (**Figure 6A)**. In the Cas12a systems, the self-targeting spacer is complementary to two or four nearby targets, all of which are positioned at intergenic sequences. The closest targets of these self-targeting sequences are adjacent to a 5’-CTTA PAM, which is recognized by conventional Cas12a nucleases (**Figure 6C**) (38, 39). There are 9 bp inverted repeats (with one mismatch) that flank Cas12a, the Rpn-family protein, and several other genes.

The Cas12a genes in these systems cover 90% of the well-characterized AsCas12a sequence (∼24% amino acid identity), including the critical crRNA-processing, DNA binding, and nuclease domains (40). Cas12a can process its own pre-crRNA via a dedicated RNAse domain (41). Three residues in this domain have been identified as critical for pre-crRNA processing; all are conserved in Rpn-associated Cas12a variants (**Figure 6D)** (41). Cas12a degrades double-stranded DNA by first cleaving the non-target strand, followed by the target strand in its single RuvC nuclease active site (42–45). Phosphate bond scission is catalyzed by two magnesium ions, one of which is coordinated by a critical aspartic residue (position 908 in *Acidaminococcus* (*As*) Cas12a). This residue is mutated to isoleucine in all Rpn-associated systems (**Figure 6D)**. Similarly, the Type I-E system encodes all the Cascade subunits but does not have Cas3. We conclude that these systems bear striking resemblance to Tn7-associated CASTs and may also mobilize genomic information for crRNA-guided horizontal transfer.

## Discussion

CASTs are rare in fully-sequenced prokaryotic genomes and are likely to be missed by traditional CRISPR detection pipelines due to their unusual operon structures and short CRISPR arrays. To address this gap, we have developed a set of Python libraries that allow users to efficiently use BLAST to search for co-occurring genes and to perform subsequent searches for arbitrary gene architectures. We have examined approximately 30 terabytes of metagenomic contigs to identify ∼1476 high-confidence CASTs, including novel Type IV and Type I-C systems, as well as a Type I-B4 CRISPR system that co-evolved with a Tn5053-like element, a member of the Tn7 family of transposons that lacks TnsA. We have also discovered systems that include a putative nuclease-inactive Cas12a and a non-Tn7 transposase-like recombinase. The diversity of CRISPR sub-types that have been co-opted by transposons will likely increase as metagenomic databases and sensitive detection pipelines continue to improve (46, 47). More broadly, the abundance of nuclease-inactive CRISPR-Cas operons may suggest that the scope of cellular processes that have co-opted CRISPR-Cas systems is wider than is currently known.

In the NCBI and metagenomic databases, the most abundant Tn7-associated CASTs are those that co-opt Class 1 CRISPR systems. Of these, Type I-F sub-systems are the most structurally diverse. Notably, we found that some I-F CASTs encode TniQ-Cas8/Cas5 fusions, duplicate Cas5s, and duplicate Cas7s. We speculate that gene duplication of the Cas5 and/or Cas7 allowed one of the paralogs to form a protein-protein interface with TniQ. The second paralog may have been subsequently lost. The remaining paralog resulted in the streamlined I-F CASTs that are most frequently found in bacterial genomes. The short atypical pre-crRNA in some I-F CASTs also suggests that these systems assemble a shortened Cascade for self-targeting. The size of Type I-E and I-F Cascades can be tuned by the length of the crRNA (48–51). Intriguingly, short I-F Cascades cannot recruit Cas3 but are still able to bind the target DNA, making them an ideal system for directed Tn7 transposition (51). Short Cascades, along with the atypical direct repeats, may differentiate self-targeting CASTs from those undergoing horizontal gene transfer in the I-F3c system.

Type I-B CASTs encode one or two TniQ/TnsD homologs. A recent report has uncovered that self-targeting in some systems proceeds via TnsD, whereas horizontal transfer is crRNA-guided (7). We also identified atypical systems that encode two short TniQ homologs along with a self-targeting spacer. Both homologs in the atypical dual-TniQ systems are related to the I-F CAST TniQ. Based on this observation, along with the crRNA-directed self-targeting, and the alignment of TniQ_3_ to the N-terminus of TniQ_1_, we propose that this CAST assembles a hetero-dimeric Cascade consisting of a single repeat of each subunit. Alternatively, this system may assemble TniQ_1_- and TniQ_3_-only Cascades for self-targeting and horizontal transfer. Additional studies will be required to decipher the role of TniQ_3_ in the atypical I-B systems with a self-targeting spacer.

How do CASTs target mobile genetic elements with minimal CRISPR arrays? We did not find any systems that retained the Cas1/Cas2 acquisition machinery, suggesting that strong evolutionary pressure is preventing the CAST-associated CRISPR arrays from expanding. CASTs encode CRISPR arrays that are significantly shorter than the corresponding canonical CRISPR-Cas systems and these arrays may also be transcriptionally silenced via *xre* elements that are frequently found adjacent to these arrays in CASTs (21). Moreover, we could not identify any CRISPR arrays in Type IV and I-C CASTs. We propose that CASTs use CRISPR arrays that occur elsewhere in the genome—perhaps in functional CRISPR-Cas systems—for horizontal gene transfer. Evidence for such *in trans* CRISPR array usage has already been documented for a large set of canonical CRISPR-Cas systems (52, 53). CRISPR arrays that are associated with active interference machinery serve as an ever-updating record of the most likely mobile genetic elements that the CAST can use for horizontal gene transfer (54). A second possibility is that Cas1/2/4 from an active CRISPR-Cas system can act *in trans* to add spacers to the CAST CRISPR array. This may be an important secondary mechanism when horizontal transfer places the CAST into a host that lacks a compatible CRISPR array.

Our search revealed Cas12a proteins associated with Rpn-family transposases, two of which appear to have atypical spacers that target two sites up- and downstream of Cas12a. This curious arrangement could be the result of a duplication of the target site that is originally present in only a single copy. We note that the HU family DNA-binding protein is only present in systems with putative self-targeting spacers and that a homolog of this protein is essential in bacteriophage Mu for transpososome assembly (55). Two other proteins of viral origin - a replisome organizer and a ribonuclease - are also found near Cas12a in a self-targeting system, hinting at an intriguing evolutionary path for the creation of this putative CAST. In sum, the CAST identification pipeline and diversity of new systems described herein add to our understanding of CRISPR transposons, expand the gene-editing toolkit, and hints at the possibility that nuclease-inactive Cas genes may play additional roles in cellular DNA metabolism.

## Materials and Methods

### Database acquisition and contig assembly

NCBI genomes were downloaded using NCBI Genome Downloading Scripts (https://github.com/kblin/ncbi-genome-download) on May 5, 2021, with the commands:

ncbi-genome-download --formats fasta bacteria

ncbi-genome-download --formats fasta archaea

Raw FASTQ files were downloaded from the EMBL-EBI repository (17) of metagenomic sequencing at ftp://ftp.ebi.ac.uk/vol1/ between January and February 2020. For each sample, the quality of the raw data was assessed with FastQC (56) using the command:

fastqc tara_reads_*.fastq.gz

Low quality reads were trimmed with Sickle (57) using the command:

sickle pe

-f name_reads_R1.fastq.gz

-r name_reads_R2.fastq.gz

-t sanger

-o name_trimmed_R1.fastq

-p name_trimmed_R2.fastq

-s /dev/null

We then used Megahit (58) to assemble the trimmed data with:

megahit

-1 name_trimmed_R1.fastq

-2 name_trimmed_R2.fastq

-o name_assembly

### Analysis pipeline

We developed a Python library, Opfi (short for Operon Finder), to search genomic or metagenomic sequence data for putative CRISPR transposons. This library consists of two modules, Gene Finder and Operon Analyzer. The Gene Finder module enables the user to use BLAST to identify genomic neighborhoods that contain specific sets of genes, such as Cas9 or TnsA. It can also identify CRISPR repeats. The Operon Analyzer module further filters the output from Gene Finder by imposing additional user-defined constraints on the initial hits. For example, Operon Analyzer can be used to find all genomic regions that contain a transposase and at least two Cas genes but no Cas3.

We used Gene Finder to locate genomic regions of interest using the following logic. First, we located all regions containing at least one transposase gene. Within those regions, we next searched for Cas genes located no more than 25 kilobase pairs away from a transposase. Transposase-containing regions without at least one nearby Cas gene were discarded from further analysis. Finally, the remaining regions were further annotated for Tn7 accessory genes (TnsC-TnsE and TniQ), and CRISPR arrays.

We further processed and categorized the Gene Finder hits using Operon Analyzer. To identify Tn7-like CRISPR-transposons, we required each putative operon to contain TnsA, TnsB, TnsC, and at least one Cas gene from Cas5-13; the distance between TnsA, TnsB, and TnsC needed to be less than 500 bp; the Cas proteins need to be downstream of TnsA/B/C and the distance between any Cas protein and TnsB needed to be less than 15 kbp. We classify this dataset into putative Class 1 systems and Class 2 systems based on their Cas signature proteins. Class 1 systems were manually reviewed to confirm the loss of adaptation (Cas1, Cas2) and interference (Cas3) proteins.

To identify non-Tn7 CRISPR-transposons, we required each putative operon to contain a CRISPR array, a transposase, and at least one Cas gene from Cas5-13. We also excluded systems containing Tn7 proteins, Cas1, or Cas2. We then partitioned this dataset into putative Class 1 systems (defined as loci with any three of Cas5/6/7/8) or Class 2 systems (Cas9, Cas12, or Cas13). For Class 1 systems, we further excluded those containing Cas3 or Cas10. To eliminate systems with fragments of effector proteins or poor matches to unrelated proteins, we required that Cas9 have a size of 2-6 kbp, Cas12 a size of 3-6 kbp, and Cas13 a size of 2.5-6 kbp. We eliminated Class 2 systems that were nucleolytically active (see below), and finally clustered all systems using mmseqs easy-cluster with a minimum sequence ID of 0.95 (59) to simplify manual curation.

### BLAST database construction

To find as many systems as possible, we assembled separate databases for Cas proteins, Tn7-family proteins, and non-Tn7 transposases. We also developed databases for common Tn7 attachment sites following a separate effort (21).

We downloaded all available bacterial and archaeal transposase sequences from UniRef50, excluding partial sequences and sequences annotated with the word “zinc” (which tended to be false positives) as well as Tn7-related proteins. All transposases associated with transposons listed in the Transposon Registry (60) were downloaded from NCBI. Finally, 100 transposases associated with each of the major families of insertion sequences were downloaded from NCBI, again excluding partial sequences, and using the ‘relevance’ sort parameter.

Amino acid sequences for Cas1–Cas13 and Tn7 family proteins (TnsA–TnsE, TniQ) were downloaded from UniRef50 (https://www.uniprot.org/uniref/). Additional Cas12 and Cas13 sequences, representing recently identified variants (e.g. Cas12k), were downloaded from the NCBI protein database (https://www.ncbi.nlm.nih.gov/protein/) and from primary literature sources (61–63).

To eliminate redundant sequences, each database was clustered using CD-HIT (http://weizhongli-lab.org/cd-hit/ (64) with a 50% sequence identity threshold and 80% alignment overlap. The clustered datasets were converted to the BLAST database format using makeblastdb (version 2.6.0 of NCBI BLAST+) with the following arguments:

makeblastdb

-in <sequence fasta file>

-title <database name>

-out <database name>

-dbtype prot

–hash_index

The full-length sequences of GuaC (PF00478), RsmJ (PF04378), YciA (PF03061) were downloaded from (http://pfam.xfam.org/). The attachment site SRP-RNA gene (ffs) (RF00169) was downloaded from RFAM (https://rfam.xfam.org/).

To assign putative Cas5-Cas8 proteins to specific Type I CRISPR-Cas subtypes, we manually collected Cas proteins and their assignments from reviews by Koonin and colleagues (62, 65, 66). All Cas protein sequences were converted into BLAST databases using makeblastdb (version 2.6.0) with default parameters.

### De-duplication of putative operons

Approximately 57% of the metagenomic systems that passed our initial filter were nearly identical at the nucleotide sequence level. However, exact nucleotide comparisons were too slow to de-duplicate this large dataset. Instead, we considered two systems to be identical if they met the following properties: (1) they had the same protein-coding genes and CRISPR arrays in the same order; (2) the genes had the same relative distances to each other; and (3) the translated sequences of all proteins were identical. This de-duplication was applied to all systems before the downstream analysis.

### Self-targeting spacer identification

Spacer sequences that were identified with PILER-CR were pairwise aligned with the contig sequence that contained them, using the Smith-Waterman local alignment function from the parasail library (67), with gap open and gap extension penalties of 8, and using the NUC44 substitution matrix. Spacers with at least 80% homology to a location in the contig were classified as self-targeting.

For Type V systems, we augmented the CRISPR array search with minCED 0.4.2 (68) after noticing transposons that were otherwise intact but seemingly lacked CRISPR arrays. The region between Cas12k and 200 bp after the end of the nearest CRISPR array was used to search for spacers (both atypical and canonical). Targets were searched for in the 500 bp region immediately downstream of the spacer search region, using the method described in the previous paragraph. For Type V systems with multiple Cas12k genes, each spacer region was aligned to each target region in order to discover systems where multiple transposons had inserted at the same attachment site.

### Phylogenetic analysis

Alignments of protein sequences were constructed with MAFFT, version v7.310 (69). Phylogenetic analysis was performed on the aligned sequences using the IQ-TREE, version 1.6.12 (70), with automatic model selection. Models used were as follows: Figure 3B: JTT+F+R3, Figure 4B cas6: PMB+G4, Figure 4B cas7: PMB+G4, Figure 4C: PMB+G4, Supplemental Figure 1 tnsB: LG+R5, Supplemental Figure 1 tnsC: LG+G4. Trees were visualized using the Figtree program version 1.4.4.

### Classification of nuclease-dead systems

To identify catalytically inactive Class 2 nucleases, we aligned each nuclease to a reference protein with MAFFT (version v7.310, with the FFT-NS-2 strategy for Cas9 and Cas12, and FFT-NS-i for Cas13). Cas9 homologs were aligned to SpCas9 (UniProtKB Q99ZW2.1, residues D10 and H840), Cas12 homologs to AsCas12a (UniProtKB U2UMQ6, residues D908 and E993), and Cas13 homologs to LbuCas13a (UniProtKB C7NBY4.1, residues R472, H477, R1048, and H1053). Mutations of D/E to anything other than D/E, or H/R to anything other than H/R/K were considered nuclease dead. As a control, we employed this technique on all 279 Cas12k proteins from NCBI as well as LbCas12a and FnCas12a (two known nuclease-active Cas12a proteins) and found that we correctly categorized them all.

### Identification of inverted repeats and target site duplications

To identify inverted repeats, we used Generic Repeat Finder (commit hash: 35b1c4d6b3f6182df02315b98851cd2a30bd6201) (71) with default parameters except as follows:

-c: 0

-s: 15

--min_tr: 15

--min_space <operon length>

--max_space <buffered operon length>

where <operon length> is the length of the putative operon and <buffered operon length> is the length of the putative operon, plus up to 1000 bp to allow a 500 bp buffer on either side of the operon. This detected inverted repeats that were at least 15 bp long. In cases where one inverted repeat fell within the bounds of the putative operon, it was discarded.

## Data availability

All processed data and analysis scripts are available on GitHub at: https://github.com/wilkelab/Metagenomics_CAST.

The analysis scripts make use of the gene_finder and operon_analyzer modules of the Opfi package, available on GitHub at: https://github.com/wilkelab/Opfi.

## Acknowledgments

We are grateful to the staff of the Texas Advanced Computing Center for providing computational resources and to members of the Finkelstein and Wilke labs for helpful discussions. This work was supported by NIGMS grants R01GM124141 (to I.J.F.) & R01GM088344 (to C.O.W.), the Welch Foundation grant F-1808 (to I.J.F.), and the College of Natural Sciences Catalyst Award for seed funding.

The authors are co-inventors on patent applications filed based on this work.

