## Supplemental Figure for "Metagenomic Discovery of CRISPR-Associated Transposons"

A

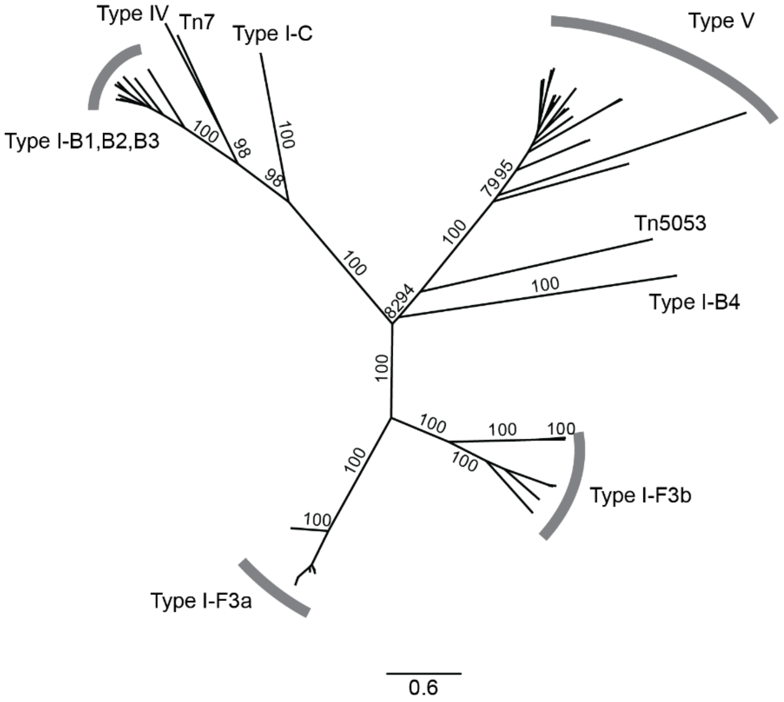


B

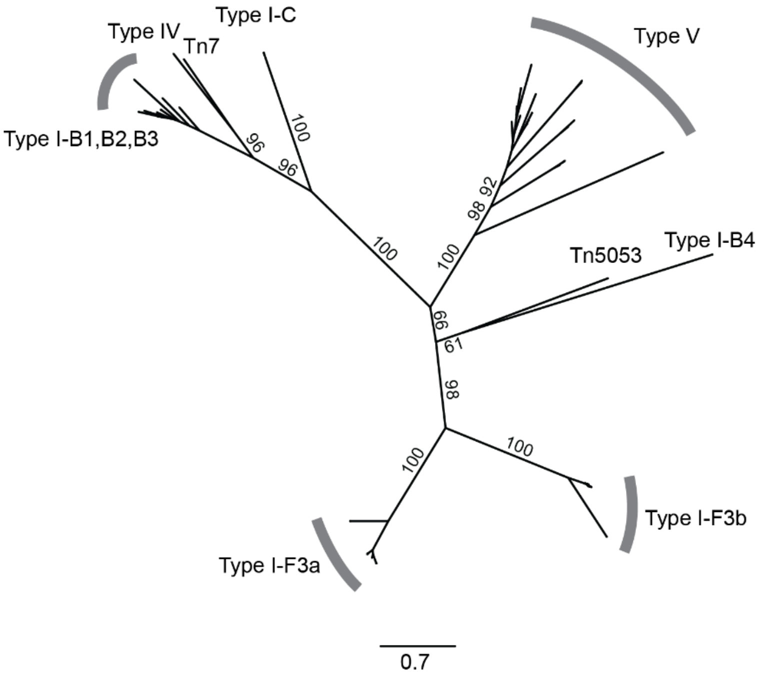


**Supplemental Figure 1**. Phylogenetic tree of (**A**) tnsB and (**B**) tnsC genes from each subtype of Tn7 CAST investigated in this work, as well as from Tn7 and Tn5053. Values at branch points are bootstrap support percentages.
